# An ultra-dense haploid genetic map for evaluating the highly fragmented genome assembly of Norway spruce *(Picea abies)*

**DOI:** 10.1101/292151

**Authors:** Carolina Bernhardsson, Amaryllis Vidalis, Xi Wang, Douglas G. Scofield, Bastian Schiffthaler, John Basion, Nathaniel R. Street, M. Rosario García-Gil, Pär K. Ingvarsson

## Abstract

Norway spruce (*Picea abies* (L.) Karst.) is a conifer species of substanital economic and ecological importance. In common with most conifers, the *P. abies* genome is very large (∼20 Gbp) and contains a high fraction of repetitive DNA. The current *P. abies* genome assembly (v1.0) covers approximately 60% of the total genome size but is highly fragmented, consisting of >10 million scaffolds. The genome annotation contains 66,632 gene models that are at least partially validated (www.congenie.org), however, the fragmented nature of the assembly means that there is currently little information available on how these genes are physically distributed over the 12 *P. abies* chromosomes. By creating an ultra-dense genetic linkage map, we anchored and ordered scaffolds into linkage groups, which complements the fine-scale information available in assembly contigs. Our ultra-dense haploid consensus genetic map consists of 21,056 markers derived from 14,336 scaffolds that contain 17,079 gene models (25.6% of the validated gene models) that we have anchored to the 12 linkage groups. We used data from three independent component maps, as well as comparisons with previously published *Picea* maps to evaluate the accuracy and marker ordering of the linkage groups. We demonstrate that approximately 3.8% of the anchored scaffolds and 1.6% of the gene models covered by the consensus map have likely assembly errors as they contain genetic markers that map to different regions within or between linkage groups. We further evaluate the utility of the genetic map for the conifer research community by using an independent data set of unrelated individuals to assess genome-wide variation in genetic diversity using the genomic regions anchored to linkage groups. The results show that our map is sufficiently dense to enable detailed evolutionary analyses across the *P. abies* genome.

## Introduction

For over a century genetic linkage maps have been used to order genetic markers and link phenotypic traits to genomic regions and chromosomes by calculating recombination events in crosses (Sturtevant 1913a; Sturtevant 1913b). With the advent of Next Generation Sequencing technologies (NGS), large numbers of markers can now be scored at a relatively low cost and within a reasonable time, which has enabled generation of high-density genetic maps consisting of thousands of markers that, in combination with a sufficiently large mapping population, can achieve unprecedented mapping resolution even in non-model systems and in species with large genomes. Genetic maps represent a complementary approach to the local, fine-scale genomic information that is available in scaffolds from a genome assembly, with a genetic map providing information on genome organization over larger scales (up to whole-chromosome level) (Fierst 2015). By grouping markers into linkage groups and subsequently ordering them within each linkage group, it is possible to anchor underlying scaffolds containing those markers to putative chromosomes with high precision (Fierst 2015). If several genetic markers, derived from a single genomic scaffold, are placed on the map, information on their relative placement in the genetic map can be used to orient the scaffold and to evaluate scaffolding decisions made in the genome assembly and hence to locate and resolve possible assembly errors (Drost et al. 2009; Bartholomé et al. 2015). For instance, when two markers originating from a single scaffold map to different linkage groups or to different regions within a linkage group, the contigs comprising the scaffold are candidates for having been wrongly joined during the assembly process. On the other hand, if markers from the same scaffold map close to each other this increases the likelihood that the scaffolding decisions were correct.

Norway Spruce (*Picea abies*) is one of the most important conifer species in Europe, both from an ecological and economic perspective. The natural distribution range of *P. abies* extends from the west coast of Norway to the Ural mountains and across the Alps, Carpathians and the Balkans in central Europe. *P. abies* composes, together with *Pinus sylvestris*, the majority of the continuous boreal forests of the Northern hemisphere where it is considered a keystone species (Farjon 1990). *P. abies* has a genome size of ∼20 Gbp that is characterized by a very high fraction of repetitive sequences. Like most conifers, *P. abies* has a karyotype consisting of 2*n*=24 and with chromosomes that are all uniformly sized (Sax and Sax 1933). Due to the large and complex genome of conifers, this important group of plants was, until recently, lacking species with available reference genomes. In 2013 the first draft assembly of the *P. abies* genome was published (Nystedt et al. 2013). Despite extensive whole-genome shotgun sequencing derived from both haploid and diploid tissues, the *P. abies* genome assembly is still highly fragmented due to the complex nature and size of the genome. The current *P. abies* genome assembly (v1.0) consists of 10.3 million scaffolds >500 bp and contains 70,736 annotated gene models of which 66,632 are at least partially validated by supporting evidence (ESTs or UniProt proteins) (Nystedt et al. 2013; De La Torre et al. 2014). Although the current genome assembly only covers about two thirds of the total genome size (12 Gbp out of the 20 Gbp *P. abies* genome), it is expected to contain the majority of expressed genes.

In this paper, we used sequence capture to identify segregating SNP markers in megagametophytes from three open-pollinated mother trees. These markers were used to create an ultra-dense haploid genetic map consisting of 21,056 probe-markers derived from 14,336 gene-bearing scaffolds in the *P. abies* genome assembly. Our aim with creating the genetic map was to 1) anchor, and where possible, order scaffolds to assign as many gene models as possible to linkage groups, and 2) to evaluate the accuracy of the *P. abies* genome assembly v1.0 on the basis of anchored scaffolds. To evaluate the accuracy of the map itself, we compared scaffold order to previously published genetic maps for *P. abies* and the closely related *Picea glauca*. Finally, we evaluated utility of the genetic map for population genomic studies by performing genome-wide analyses of genetic diversity for the genomic regions anchored in the map using a sample of c. 500 unrelated *P. abies* trees.

## Material and Methods

### DNA extraction and sequence capture

In the autumn of 2013, seeds were collected for linkage map construction from five of 30 putative ramets of Z4006, the genotype used to generate the reference genome for *Picea abies* (Nystedt et al. 2013). Megagametophytes were dissected from 2,000 seeds by removing the diploid seed coat surrounding the haploid megagametophyte tissue. DNA extraction from megagametophytes was performed using a Qiagen Plant Mini Kit. Each extracted sample was measured for DNA quality using a Qubit^®^ ds DNA Broad Range (BR) Assay Kit, and all samples with a total amount of DNA >354 ng were kept. The remaining 1,997 samples were sent to RAPiD Genomics© (Gainesville, Florida, USA) in September 2014 for sequence capture using 31,277 capture probes that had been specifically designed to target 19,268 partially-validated gene models from the *P. abies* genome assembly. Where possible, probes were designed to flank regions of known contig joins in the v1.0 genome assembly (for further detail of the probe design, see Vidalis et al. 2018).

The capture data was sequenced by RAPiD Genomics© on an Illumina HiSeq 2000 using 1×75 bp sequencing and was delivered in October 2015. The raw reads were mapped against the complete *P. abies* reference genome v.1.0 using BWAMEM v.0.7.12 and default settings (Li and Durbin 2009). Following read mapping, the genome was subset to only contain the probe-bearing scaffolds (a total of 18,461 scaffolds) using Samtools v.1.2 (Li and Durbin 2009; Li et al. 2009). Duplicates were marked and local realignment around insertion/deletions (indels) was performed using Picard (http://broadinstitute.github.io/picard) and GATK (https://software.broadinstitute.org/gatk) (McKenna et al. 2010; DePristo et al. 2011). Genotyping was performed using GATK Haplotypecaller (version 3.4-46, (DePristo et al. 2011; Van der Auwera et al. 2013) with a diploid ploidy setting and gVCF output format. We used a diploid ploidy setting to increase the likelihood of detecting possible sample contamination from diploid tissue for the haploid megagametophyte samples. CombineGVCFs was then run on batches of ∼200 gVCFs to hierarchically merge them into a single gVCF and a final SNP call was performed using GenotypeGVCFs jointly on the 10 combined gVCF files, using default read mapping filters, a standard minimum confidence threshold for emitting (stand-emit-conf) of 10, and a standard minimum confidence threshold for calling (stand_call_conf) of 20. See Vidalis et al. (2018) and the script “per_sample_gvcf.sh” (available at https://github.com/parkingvarsson/HaploidSpruceMap) for a full description of the pipeline used for calling variants.

### SNP filtration and megagametophyte relationships

After SNP filtering, we performed a principle component analysis (PCA) to evaluate the relationship among samples (see Supplementary file for details on the PCA analysis and subsequent filtering steps). Based on the PCA and a hierarchical clustering approach, we divided samples into three clusters representing putative maternal families (Supplementary, Figure S1-3) that were then analyzed independently. In the end we obtained 9,073 probe-markers from 7,101 scaffolds for Cluster 1 (314 samples), 11,648 probe-markers from 8,738 scaffolds for Cluster 2 (270 samples) and 19,006 probe-markers from 13,301 scaffolds for Cluster 3 (842 samples) with a total of 21,056 probe-markers from 14,336 scaffolds across all three clusters (Table 1). In total, these scaffolds cover 0.34 Gbp of the *P. abies* genome and contain 17,079 partially validated gene models.

**Table 1:** Overview of the three component maps and the total number of probe-markers available in the consensus map. Cluster: Name of each putative maternal family that was identified in the principal component analysis. Samples: Number of megagametophytes in each cluster. Markers: Number of probe-markers in each component map with number of unique segregating bins within brackets (one marker for each bin was used to anchor the bin markers to the genetic map). Scaffolds: Number of scaffolds represented in each component map.

| Cluster | Samples | Markers | Scaffolds |
| --- | --- | --- | --- |
| Cluster 1 | 314 | 9,073 (3,924) | 7,101 |
| Cluster 2 | 270 | 11,647 (5,311) | 8,738 |
| Cluster 3 | 842 | 19,006 (11,479) | 13,301 |
| Total | 1,426 | 21,056 | 14,336 |

### Component and consensus maps

We created genetic linkage maps using the R-package BatchMap (Schiffthaler et al. 2017), a parallel implementation of the R-package Onemap (Margarido, Souza, and Garcia 2007). All probe-markers were recoded using the D1.11 cross-type (Wu et al. 2002), tested for segregation distortion (p < 0.05 after Bonferroni correction) (Supplementary, Figure S4) and grouped into marker bins. The probe-marker with lowest amount of missing data in each bin was then used to represent the bin when constructing the genetic map. Bin markers were grouped into LGs using LOD = 8 and a maximum recombination fraction = 0.35. LGs were then ordered using the RECORD algorithm (Van Os et al. 2005) with 16 times counting, parallelized over 16 cores, reordered in a 10 marker sliding window with 1 marker incremental steps using the command ‘ripple’ and finally mapped using the Kosambi mapping function and the ‘map batches’ approach (Schiffthaler et al. 2017) over four parallel cores. Finally, heat maps with pairwise recombination fraction (lower triangular) and phase LOD score (upper triangular) for the ordered markers were created to evaluate the ordering accuracy of independent linkage groups (Supplementary, Figure S5 and S6A-L). We observed 183 probe-marker bins showing signs of segregation distortion. These bins were, however, randomly distributed over the linkage groups and did not appear to affect marker ordering and map distance and were therefore retained in subsequent analyses.

To evaluate correspondence between LGs in maps derived from the three PCA clusters, the number of unique scaffolds shared between cluster LGs were counted (Supplementary, Figure S5). We then created a consensus map for each linkage group from the three independent component maps using the R-package LPmerge (Endelman and Plomion 2014) with component maps ranked according to marker numbers (Cluster 3, Cluster 2, Cluster 1), a maximum interval setting ranging from one to 10 and map weights proportional to the size of the mapping population (Cluster 3= 0.5, Cluster 2 and Cluster 1 = 0.25). From all possible consensus maps generated by LP merge, for each linkage group we selected the map with the lowest mean root mean square error (RMSE) to serve as the consensus map (Endelman and Plomion 2014). Order correlations between individual component maps and the consensus maps (Table 2 and Supplementary, Figure S7A-L) as well as between the three component maps (Supplementary, Figure S8A-L) were estimated using Kendall’s τ. For visual representation of the consensus map we created a Circos plot using the R-package omicCircos (Hu et al. 2014), available from Bioconductor (https://bioconductor.org/biocLite.R).

**Table 2:** Marker density and size of each component genetic map created from the three clusters as well as for the consensus map. LG: Linkage group. Cluster 1-3: Component maps for cluster 1-3 with number of probe-markers (marker-bins) assigned, map size (in cM) and maximum gap in map (in cM) for each of the LGs. Consensus: Number of markers and map size of the LGs in the consensus map.

| LG | Cluster 1 |  |  | Cluster 2 |  |  | Cluster 3 |  |  | Consensus |  |
| --- | --- | --- | --- | --- | --- | --- | --- | --- | --- | --- | --- |
|  | Markers | Length (cM) | Max gap (cM) | Markers | Length (cM) | Max gap (cM) | Markers | Length (cM) | Max gap (cM) | Markers | Length (cM) |
| I | 975 (421) | 385.5 | 8.0 | 1,159 (553) | 439.9 | 21.1 | 1,967 (1,185) | 414.1 | 8.8 | 2,172 | 414.1 |
| II | 701 (305) | 249.2 | 9.6 | 863 (366) | 289.0 | 9.4 | 1,456 (864) | 289.8 | 10.9 | 1,608 | 250.3 |
| III | 859 (394) | 324.0 | 4.6 | 1,069 (479) | 381.1 | 7.1 | 1,738 (1,075) | 346.4 | 5.2 | 1,940 | 342.5 |
| IV | 771 (323) | 298.7 | 14.5 | 970 (452) | 350.9 | 8.6 | 1,531 (916) | 303.0 | 27.0 | 1,704 | 303.0 |
| V | 761 (311) | 273.2 | 8.9 | 1,116 (499) | 395.6 | 9.5 | 1,649 (1,032) | 342.6 | 15.1 | 1,865 | 275.0 |
| VI | 648 (292) | 241.0 | 8.4 | 915 (399) | 270.7 | 4.6 | 1,456 (894) | 269.5 | 8.4 | 1,622 | 240.2 |
| VII | 682 (331) | 314.0 | 8.4 | 923 (443) | 380.8 | 13.4 | 1,625 (1,013) | 321.9 | 7.9 | 1,769 | 321.0 |
| VIII | 775 (339) | 307.0 | 5.6 | 943 (454) | 367.26 | 9.8 | 1,465 (904) | 315.6 | 6.6 | 1,609 | 305.9 |
| IX | 792 (332) | 283.3 | 5.4 | 786 (364) | 295.6 | 5.9 | 1,589 (911) | 285.1 | 7.4 | 1,738 | 285.0 |
| X | 648 (289) | 231.6 | 7.0 | 960 (454) | 342.7 | 6.9 | 1,564 (917) | 272.7 | 7.1 | 1,709 | 273.1 |
| XI | 677 (253) | 200.6 | 3.7 | 1,025 (411) | 269.2 | 4.0 | 1,440 (818) | 233.6 | 3.0 | 1,608 | 233.4 |
| XII | 784 (334) | 281.6 | 9.3 | 919 (437) | 360.7 | 11.1 | 1,526 (950) | 312.3 | 14.3 | 1,712 | 312.3 |
| Total | 9,073 (3,924) | 3,389.4 | 14.5 | 11,648 (5,311) | 4,143.4 | 21.1 | 19,006 (11,479) | 3,706.7 | 27.0 | 21,056 | 3,555.8 |

To evaluate the inflation of map distances due to possible genotyping errors, we performed 100 rounds of random subsampling of 100 probe-marker bins per LG and component map. The following marker ordering and genetic distance calculation were performed with 10 rounds of RECORD and the Kosambi mapping function.

### Accuracy of the reference Picea abies genome assembly

To evaluate the accuracy of scaffolds from the v1.0 *P. abies* reference genome containing at least two probe-markers (here after called multi-marker scaffolds) we determine whether probe-markers from the same genomic scaffold mapped to the same region of an LG, on different regions within a single LG or on different LGs. In the consensus map, we considered markers to be positioned in the same region on an LG if all probe-markers from a scaffold mapped within a 5 cM interval of each other. If any marker from the scaffold was positioned further apart, the scaffold was tagged as containing a putative assembly error. The same considerations were made for scaffolds with probe-markers positioned on different LGs.

### Comparative analyses of Picea linkage maps

To evaluate the consistency of our genetic map with earlier maps from *P. abies* we compared our haploid consensus map to the *P. abies* linkage map from Lind et al. (2014). The Lind et al. map was created using genetic markers generated using an Illumina 3072 SNP Golden Gate Assay. We performed using `tblastn` sequence homology searches against the *P. abies* v1.0 genome assembly for the SNP array sequences of the makers mapped in the Lind et al. map and extracted reciprocal best hits with >95% identity, which were then assigned to the corresponding scaffold in the *P. abies* genome. We performed similar analyses to compare the synteny between our consensus map and the *P. glauca* composite map from Pavy et al. (2017). Again, we used `tblastn` sequence homology search comparisons of array sequences from the *P. glauca* SNP array (Pavy et al. 2013) with scaffolds from the *P. abies* v1.0 genome assembly to assign corresponding map positions between *P. abies* and *P. glauca*. In order to evaluate correspondence between LGs from the different genetic maps, we assessed the number of shared scaffolds between our consensus map, the Lind et al. and Pavy et al. maps. Consistency of scaffold ordering was then evaluated using visual comparisons (Figure 4 and 5) and by calculating correlations of marker orders using Kendall’s τ.

### Population genetic analysis of the consensus genetic map

In order to independently evaluate the utility of the consensus map for downstream research, we used a subset of the data from Baison et al. (2018) to estimate patterns of nucleotide diversity across the Norway spruce genome. The data from Baison et al. originally contained 517 individuals sequenced with 40,018 probes designed for diploid spruce samples (Vidalis et al 2018). We extracted data for all probes that we had anchored in our genetic map from the VCF file containing the data from Baison et al.. We further hard-filtered the resulting VCF file by only considering bi-allelic SNPs within the extended probe regions (120 bp probes ±100 bp) with a QD >5, MQ >50 and a overall DP between 3000 and 16000. Samples containing >25% missing data were removed from further analysis. We used the data to calculate nucleotide diversity (π), the number of segregating sites and Tajima’s D (Tajima 1989). We used the R package vcfR (Knaus and Grünwald 2017) to read the VCF-file into R and then used in-house developed scripts to perform all calculations (available at https://github.com/parkingvarsson/HaploidSpruceMap). We assigned probes to LGs and map positions by assigning them the coordinates of the physically closest (in bp) probe. We also calculated pairwise linkage disequilibrium (LD) between markers within probes using vcftools (Danecek et al. 2011) and imported the results into R where they were used to calculate *Z_nS_* scores (Kelly 1997) per probe using an in-house developed script (available at https://github.com/parkingvarsson/HaploidSpruceMap). Finally, we ran sliding window analyses along the linkage groups for the different summary statistics using 10 cM windows that were moved in 1 cM incremental steps.

## Results

We generated a *P. abies* consensus linkage map from three haploid component maps containing a total of 21,056 unique probe-markers from 14,336 scaffolds in the *P. abies* genome assembly v1.0. The consensus map anchored 0.34 Gbp of the *P. abies* 1.0 assembly, corresponding to 1.7% of the complete *P. abies* genome or 2.8% of the genome assembly. However, these scaffolds anchor 25.6% of all validated gene models with these anchored scaffolds containing 31.7%, 20.6% and 25.8% of the High‐, Medium‐ and Low confidence gene models from Nystedt et al (2013), respectively. The consensus map had a total length of 3,556 centiMorgan (cM), distributed over 12 linkage groups (LGs), corresponding to the haploid chromosome number (Sax and Sax 1933), and with an average distance of 0.17 cM between probe-markers (Table 2, Figure 1A).

**Figure 1:**
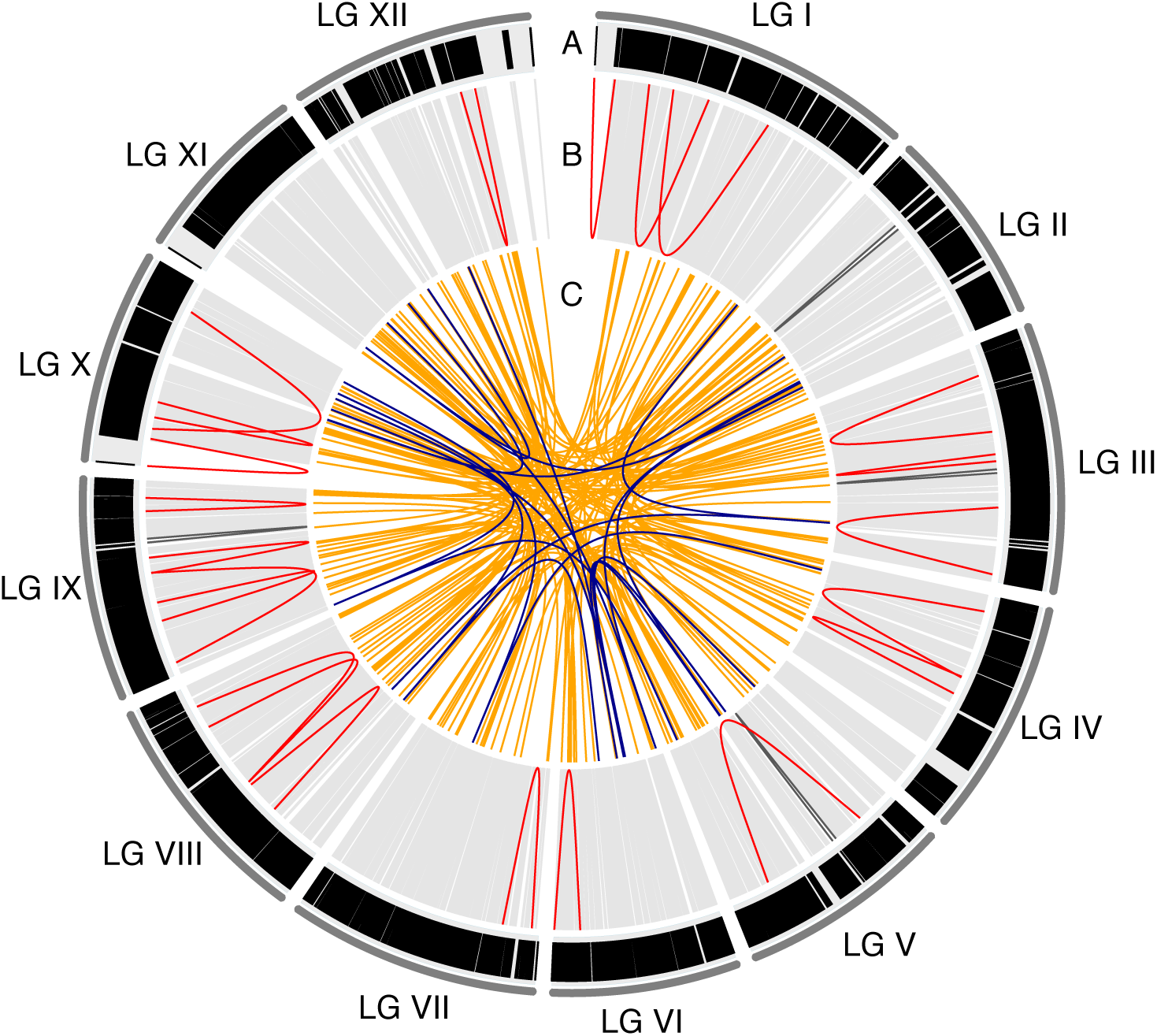
Circos plot of the consensus map. A) Marker distribution over the 12 linkage groups (LG I-LG XII). Each black vertical line represents a marker (21,056 in total) in the map and is displayed according to the marker positions in cM. Track B-C visualizes multi marker scaffolds, where each line is a pairwise position comparison of probe-markers from the same scaffold. B) Position comparisons of probe-markers from the same scaffold that are located on the same LG. Light grey lines indicate probe-markers that are located < 5cM from each other, dark grey lines indicate probe-markers located 5-10 cM apart and red lines indicate probe-markers >10 cM apart. C) Position comparisons of probe-markers from the same scaffold that are mapping to different LGs. Orange lines indicated probe-markers from the same scaffold split over 2 LGs, while dark blue lines indicated probe-markers split over 3 LGs.

Correlations of probe-marker order between the three component maps and the consensus map ranged from 0.96 to 0.998, while the correlations between marker orders between individual component maps ranged from 0.943 to 0.993 (Supplementary, Figure S7 and S8). 183 probe-marker bins showed evidence of segregation distortion in Cluster 3, but these were randomly distributed over all linkage groups and we did not observe regions showing clusters of markers with segregation distortion or with conflicting marker orders between clusters (Supplementary, Figure S8). LG XI, which displayed the largest discrepancy in marker order between component maps, has a region at the distal end of the LG, covering 252 probe-markers, where the resolution was too low to identify the correct marker order and where the entire region was positioned at 36.115 cM (Supplementary, Figure S7K and S8K), explaining the lower correlations in marker order between individual maps for this LG.

We used a random subsampling approach to evaluate potential inflation of map distances due to possible genotyping errors. From these analyses, total map size for Cluster 1 ranged between 2,166.8 and 2,450.0 cM with an average size of 2,294.2 cM and a standard deviation (SD) of 3.6- 5.8 cM per LG. Cluster 2 ranged between 2,304.2 and 2,663.6 cM with an average of 2,478.3 cM and a SD of 4.4 – 9.1 cM per LG, while Cluster 3 ranged between 1,855.4 and 2,093.2 cM with an average of 1,971.0 cM and a SD of 2.7 – 7.3 cM per LG. The estimated inflation was therefore predicted to be 0.15 – 0.31 cM per probe-marker bin across the three component maps (Table 3). This inflation per probe-marker bin roughly corresponded to the map resolution of the clusters (Cluster 1 - 0.32 cM: Cluster 2 - 0.37 cM: Cluster 3 – 0.12 cM) and yielded an error estimate of ∼1 genotype error per marker-bin or 11-17 genotype errors per sample.

**Table 3:** Estimated genetic length of each Linkage Group (LG) in the three component maps. LG: linkage group in the consensus map; Observed genetic length (cM): The genetic length of the LG calculated from all probe-marker bins (same as in table 2); Mean estimated genetic length (cM): the average length of the LG when using 100 random probe-marker bins in 100 map calculations; SD (cM): Standard deviation of the estimated length; Inflation/Marker bin: The difference between observed genetic length and the estimated length divided by the number of probe-marker bins in the linkage group.

| LG | Cluster 1 |  |  |  | Cluster 2 |  |  |  | Cluster 3 |  |  |  |
| --- | --- | --- | --- | --- | --- | --- | --- | --- | --- | --- | --- | --- |
|  | Observed<br>genetic<br>length<br>(cM) | Mean<br>estimated<br>genetic<br>length<br>(cM) | SD<br>(cM) | Inflation /<br>Marker bin<br>(cM) | Observed<br>genetic<br>length<br>(cM) | Mean<br>estimate<br>d genetic<br>length<br>(cM) | SD<br>(cM) | Inflation /<br>Marker bin<br>(cM) | Observed<br>genetic<br>length<br>(cM) | Mean<br>estimated<br>genetic<br>length<br>(cM) | SD<br>(cM) | Inflation /<br>Marker bin<br>(cM) |
| I | 385.5 | 245.5 | 5.2 | 0.33 | 439.9 | 252.3 | 6.1 | 0.34 | 414.2 | 204.8 | 7.3 | 0.18 |
| II | 249.2 | 168.8 | 4.7 | 0.26 | 289.0 | 192.9 | 4.4 | 0.26 | 289.8 | 166.4 | 2.8 | 0.14 |
| III | 324.0 | 195.8 | 5.8 | 0.33 | 381.1 | 218.6 | 5.8 | 0.34 | 346.4 | 168.5 | 3.9 | 0.17 |
| IV | 298.7 | 204.7 | 5.0 | 0.29 | 350.9 | 215.6 | 5.7 | 0.30 | 303.0 | 167.0 | 3.5 | 0.15 |
| V | 273.2 | 195.7 | 4.6 | 0.25 | 395.6 | 218.4 | 9.0 | 0.36 | 342.6 | 180.0 | 5.1 | 0.16 |
| VI | 241.0 | 161.8 | 4.7 | 0.27 | 270.7 | 170.0 | 4.7 | 0.25 | 269.5 | 142.2 | 2.9 | 0.14 |
| VII | 314.0 | 223.6 | 5.3 | 0.27 | 380.8 | 248.7 | 6.3 | 0.30 | 321.9 | 175.9 | 3.7 | 0.14 |
| VIII | 307.0 | 203.6 | 4.9 | 0.31 | 367.26 | 226.7 | 5.7 | 0.31 | 315.6 | 179.2 | 4.3 | 0.15 |
| IX | 283.3 | 194.0 | 4.9 | 0.27 | 295.6 | 185.3 | 6.8 | 0.30 | 285.1 | 157.2 | 3.0 | 0.14 |
| X | 231.6 | 164.4 | 3.6 | 0.23 | 342.7 | 193.6 | 4.5 | 0.33 | 272.7 | 141.5 | 2.7 | 0.14 |
| XI | 200.6 | 141.8 | 4.7 | 0.23 | 269.2 | 147.0 | 4.8 | 0.30 | 233.6 | 119.6 | 3.0 | 0.14 |
| XII | 281.6 | 194.4 | 4.9 | 0.26 | 360.7 | 209.6 | 9.1 | 0.35 | 312.3 | 168.7 | 3.1 | 0.15 |
| Total | 3,389.4 | 2,294.2 | - | 0.28 | 4,143.4 | 2,478.3 | - | 0.31 | 3,706.7 | 1,971.0 | - | 0.15 |

### Evaluation of the Picea abies genome assembly v1.0

4,859 scaffolds (33.9%) contained more than one unique probe-marker combined over all three component maps. 185 of these multi-marker scaffolds contained markers that were located in more than one LG (*inter-split scaffolds*) or over different parts of the same LG (*intra-split scaffolds*). 26 scaffolds (0.18% of mapped scaffolds and 0.54% of multi-marker scaffolds) contained markers that were positioned on the same LG but at distances exceeding 5 cM in the consensus map. When exploring the individual component maps, it was apparent that for two of these scaffolds (MA_281725 on LG X and MA_10431182 on LG I) the probe-markers in the consensus map all came from different component maps The consensus map thus contain a gap that we can not verify using any of the individual component maps (Figure 1 and Supplementary, Figure S9). Three other scaffolds (MA_9458 on LG IX, MA_10431315 on LG II and MA_10432328 on LG III) all have multiple probe-markers present in at least one component map and were these component maps do not support the split we observe in the consensus map (Supplementary, Figure S9). It thus appears that these splits are artifacts arising from the construction of the consensus map.

There were 164 scaffolds (1.14% of mapped scaffolds and 3.38% of multimarker scaffolds) containing markers that were mapped to two or three different LGs (Figure 2 and Supplementary, Figure S10). All LGs contained inter-split scaffolds, while 10 LGs (LGII and LGXI are the exceptions) contained intra-split scaffolds supported by the component maps (Figure 1B-C and Supplementary, Figure S9).

**Figure 2:**
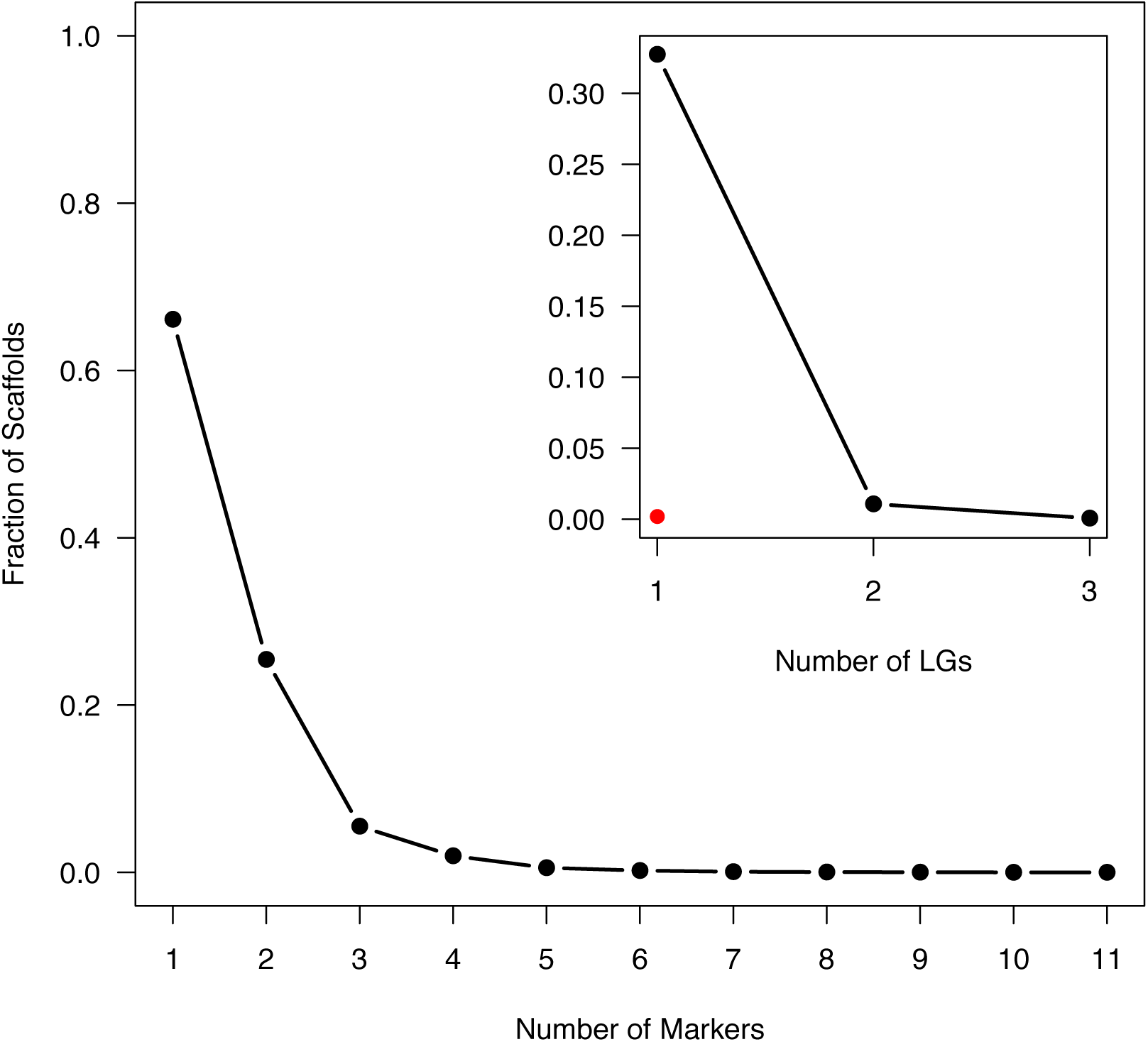
Fraction of scaffolds that are being represented by 1-11 unique markers in the consensus map. Insert: Fraction of scaffolds that have multiple probe-markers (2-11) that are distributed over 1-3 linkage groups (inter-split scaffolds). Red dot indicate the fraction of scaffolds with multiple probe-markers which are positioned > 5cM apart on the same linkage group (intra-split scaffolds).

The scaffolds covered by the consensus map ranged in length from 0.22 to 208.1 Kbp with a median of 17.1 Kbp, while multi-marker scaffolds ranged from 0.39 to 161.5 Kbp (median of 21 Kbp). The 185 scaffolds that are split within or across LGs ranged in size from 2.5 to 121.6 Kbp, with a median length of 36.9 Kbp. Split scaffolds were significantly longer than multi-marker scaffolds in general (t = −7.7, df = 193.4, p-value = 7.0e-13; Figure 3), suggesting that longer scaffolds are more likely to contain assembly errors compared to shorter scaffolds. Split scaffolds mostly contained high‐ and medium confidence gene models (Table 4). A visual inspection of the split scaffolds revealed that for 75 and 10 of the inter-split and intra-split scaffolds, respectively, the predicted position of the split(s) occurred between different gene models on the same scaffold. Of greater concern, for 88 of the inter-split scaffolds and 11 of the intra-split scaffolds the predicted position of the split was located within a single gene model (Supplementary, Figure S9 and S10). In addition, 21 inter-split scaffolds showed an even more complicated picture, where an interior region of the gene model (most often containing an intron > 5kb) mapped to another chromosome whereas the 5’ and 3’ regions of the gene model mapped to the same chromosome location (Supplementary, Figure S10). However, 84% (184 out of a total of 219 splits) appear to occur between contig joins (where a sequence of N’s appear in the assembly) of the scaffold. Of the 17,079 gene models that were anchored to the consensus genetic map, 330 were positioned on inter‐ or intra-split scaffolds (5.4% of gene models that were positioned on multi-marker scaffolds) and 100 showed a split within gene models (1.6% of gene models from multi-marker scaffolds) (Table 4).

**Figure 3:**
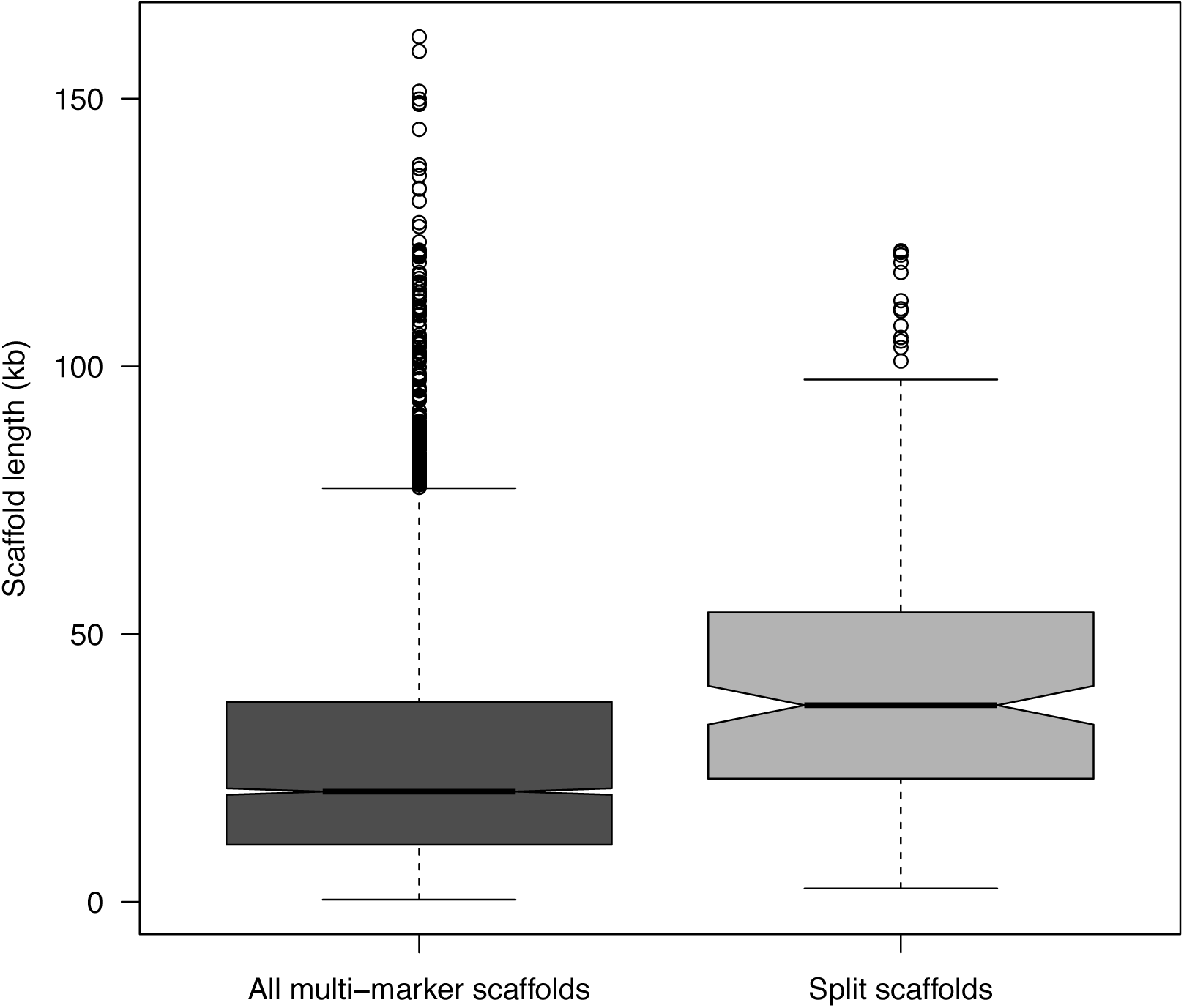
Box plot of scaffold lengths for all multi-marker scaffolds (dark gray box) and for scaffolds showing a split within or across LGs (light gray box). The split scaffolds are significantly longer than the multi-marker scaffolds in general (t = −7.70, df = 193.39, p-value = 7.00e-13).

**Table 4:** Overview of annotated gene models anchored to the genetic map. Gene models: Annotated protein coding gene models with High-, Medium‐ and Low confidence level (Nystedt et al. 2013). Mapped scaffolds: Number of gene models positioned on scaffolds that are anchored to the genetic map (Percentage of total number of gene models for each confidence level). Multi-marker scaffolds: Number of gene models positioned on scaffolds with multiple markers in the genetic map (Percentage of gene models on mapped scaffolds). Inter-split scaffolds: Number of gene models positioned on the 164 scaffolds that are split between LGs in the genetic map (Percentage of gene models on mapped scaffolds / Percentage of gene models on multi-marker scaffolds). Intra-split scaffolds: Number of gene models positioned on the 22 scaffolds that are split between different regions of the same LG (Percentage of gene models on mapped scaffolds / Percentage of gene models on multi-marker scaffolds). Split within gene models: Number of gene models that have an internal split (Percentage of gene models on mapped scaffolds / Percentage of gene models on multimarker scaffolds).

| Gene models | Mapped scaffolds | Multi-marker scaffolds | Inter-split scaffolds | Intra-split scaffolds | Split within gene models |
| --- | --- | --- | --- | --- | --- |
| High confidence | 8,379 (31.7%) | 3,122 (37.3%) | 145 (1.7% / 4.6%) | 15 (0.18% / 0.48%) | 58 (0.69% / 1.9%) |
| Medium confidence | 6,624 (20.6%) | 2,215 (33.4%) | 114 (1.7% / 5.1%) | 16 (0.23% / 0.68%) | 29 (0.44% / 1.3%) |
| Low confidence | 2,076 (25.8%) | 762 (36.7%) | 35 (1.7% / 4.6%) | 5 (0.29% / 0.79%) | 13 (0.63% / 1.7%) |
| Total | 17,079 (25.6%) | 6,099 (35.7%) | 294 (1.7% / 4.8%) | 36 (0.21% / 0.59%) | 100 (0.59% / 1.6%) |

### Comparative analyses to other Picea linkage maps

In order to assess the accuracy and repeatability of the *P. abies* genetic maps we compared our consensus map to the *P. abies* map presented in Lind et al. (2014). 353 comparisons between 298 markers from Lind et al. and 288 scaffolds contained in our consensus map were identified at a > 95 % identity threshold. Of these markers, 96.7% grouped to the same LG in the two maps while the remaining 3.3% (11 out of 353) were distributed across several LGs (Figure 4). Correlations of marker order between the two *P. abies* maps ranged from 0.53 to 0.99 across the 12 LGs. The comparison between the haploid consensus map for LG I and LG 7 from Lind et.al, which had the lowest correlation of marker order, showed inconsistencies of marker order where a contiguous subset of markers were arranged in the opposite order from the rest of the markers for that LG. The remaining LGs showed high synteny, with consistent marker ordering between the two genetic maps.

**Figure 4:**
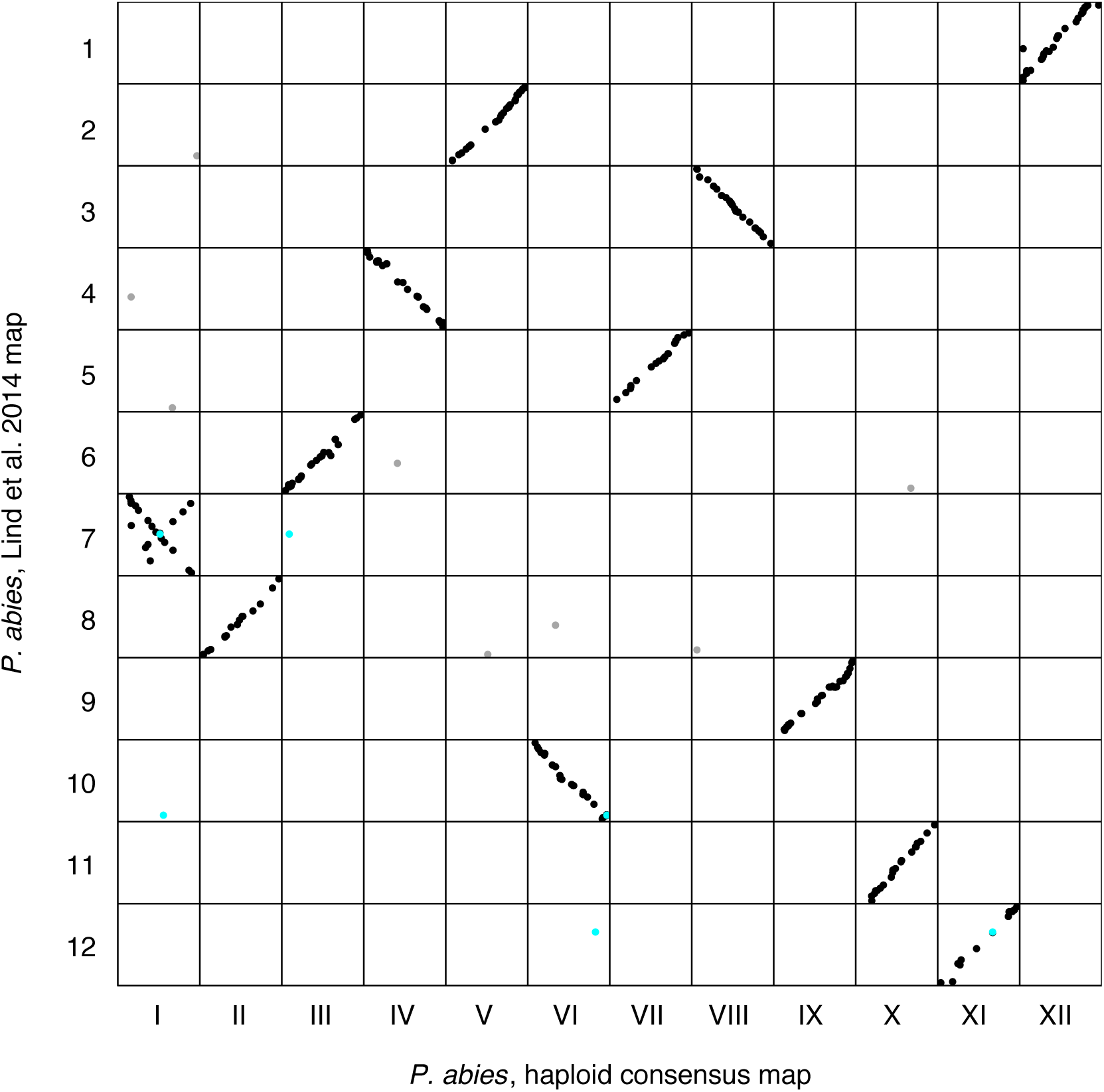
Marker order comparison between Linkage Groups (LGs) from the haploid consensus map presented here and the *Picea abies* map from Lind et al. (2014). Consensus LG I - LG XII are located on the x-axis from left to right. Lind et al. LG 1 -LG 12 are located on the y-axis from top to bottom. Each dot represents a marker comparison from the same scaffold, where black coloration represents the LG where the majority of marker comparisons are mapped. Grey coloration represents markers mapping to a different LG compared to the majority of markers. Turquoise coloration represents markers located on split scaffolds, which are indicative of assembly errors.

Synteny between *P. abies* and *P. glauca* species was assessed by comparing LG location and marker order between our *P. abies* consensus map and the composite map of *P. glauca* from Pavy et al. (2017). 14,112 comparisons of 4,053 gene models in the composite map in *P. glauca* (Pavy et al. 2017) and 4,310 scaffolds in the *P. abies* consensus map were identified at a > 95% identity threshold. 92.7% (13,084 out of 14,112 comparisons) of these were located on homologous LGs while the remaining 7.3% (1,028 comparisons from 388 *P.abies* scaffolds) were distributed across the 12 LGs (Figure 5). 8.2% of all comparisons from multi-probe scaffolds were between non-homologous LGs while 44.3% of all comparisons from split scaffolds were between non-homologous LGs. 31.9% of all non-homologous LG comparisons involved split scaffolds. The correlations of marker order between the two maps were comparable to the correlations we observed between individual component maps in *P. abies* (0.96-0.99), showing that synteny is highly conserved between *P. abies* and *P glauca*.

**Figure 5:**
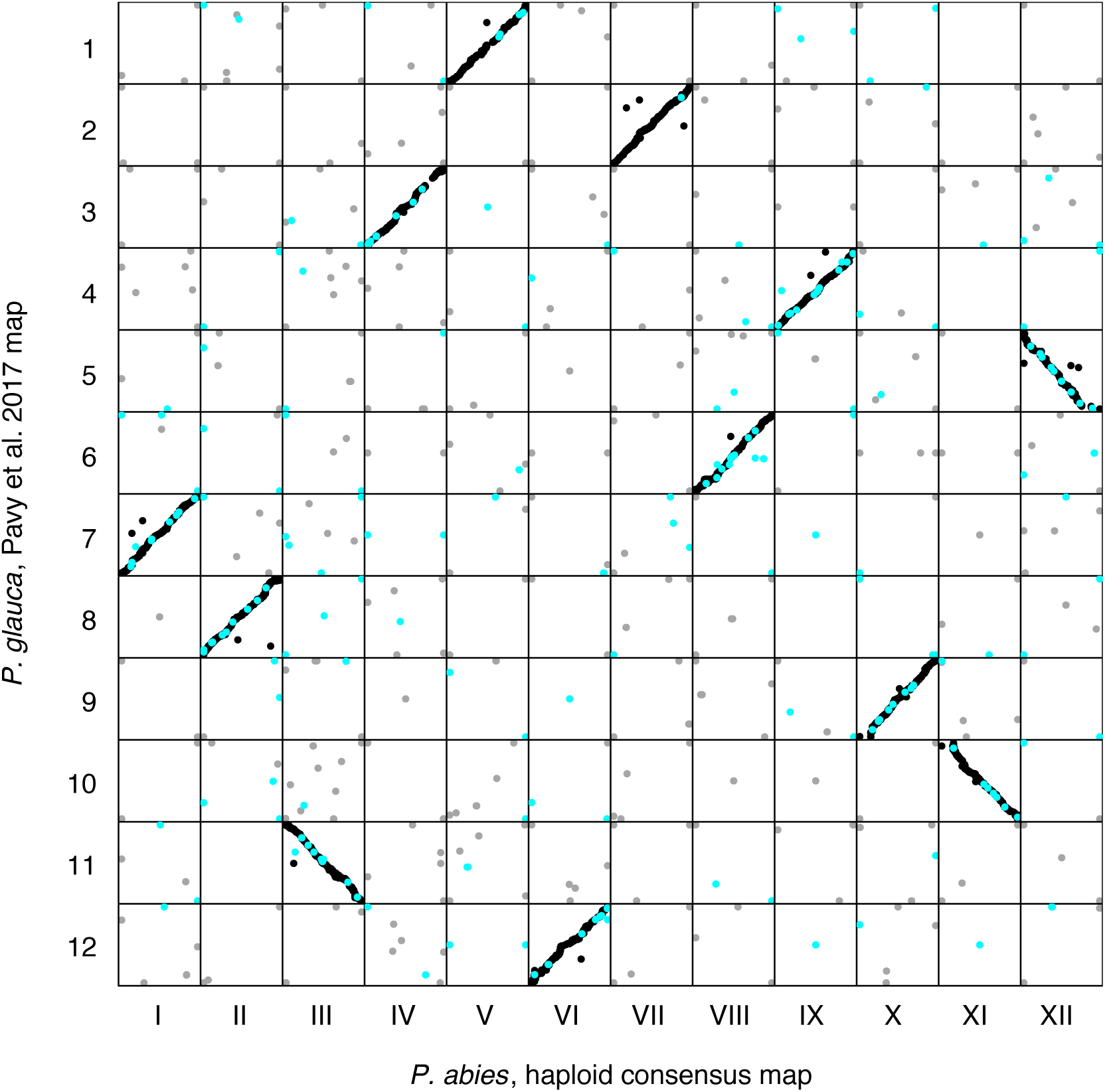
Marker order comparison of Linkage Groups (LGs) between the *Picea abies* haploid consensus map presented here and the *Picea glauca* map from Pavy et al. (2017). Consensus LG I - LG XII are located on the x-axis from left to right. Pavy et al. LG 1 - LG 12 are located on the y-axis from top to bottom. Each dot represents a marker comparison from the same scaffold, where black coloration represents markers mapping to the same LG in the two species, grey coloration represents markers mapping to different LGs. Turquoise coloration represents markers located on split scaffolds, indicating an assembly error.

### Population genetic analyses based on the consensus map

22,413 probes, covering 12,908 scaffolds, were used in the population genetic analyses based on the consensus genetic map. On a per-probe basis, we observed substantial variation in all neutrality statistics, with the number of segregating sites ranging from 0 - 77 (mean 15.9), nucleotide diversity (π) from 0 - 0.4 (0.005), Zns from 0 - 1 (mean 0.04) and Tajima’s D from −2.4 – 3.5 (mean −0.85). To study large-scale trends and possible chromosomal differences we performed sliding window analyses across the LGs for the different summaries (Figure 6). One interesting large-scale feature we observed was that SNP densities were often highest at the distal or central regions of LGs, indicating the possible location of centromeres and telomeres, for which recombination rates are expected to be reduced (Gaut et al. 2007) and where we hence would expect higher densities of probes per cM (Figure 6a). The large-scale analyses also revealed several instances where entire chromosomal arms might be experiencing different evolutionary patterns (Figure 6b-c). Finally, we identified regions that appear to be evolving under the influence of natural selection. For instance, several regions showed higher than average levels of nucleotide diversity and positive Tajima’s D (e.g. on LG IV, V and XII), suggesting that they might harbor genes under balancing selection. Similarly, regions with low nucleotide diversity, an excess of rare alleles and strong linkage disequilibrium (i.e. negative Tajima’s D and high *Z_ns_* scores, e.g. on LG III) could indicate regions harboring possible selective sweeps (Figure 6c-d).

**Figure 6:**
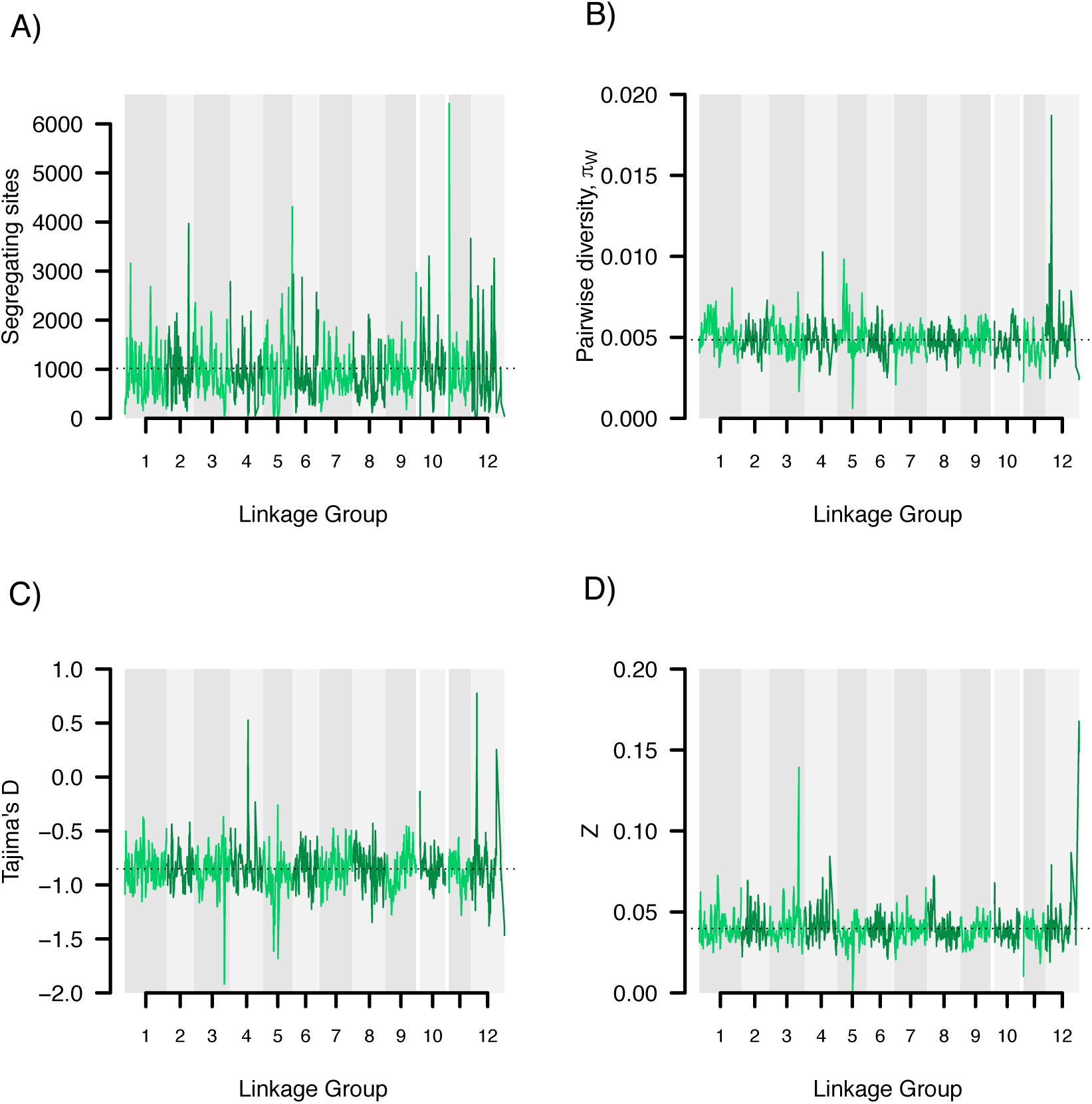
Sliding window analysis of neutrality statistics. Analyses were performed using 10 cM windows with 1 cM incremental steps along the consensus map linkage groups and visualized using coloring alternates between adjacent LGs. A) Number of segregating sites. Dashed horizontal line indicates the overall average of 1017. B) Pairwise nucleotide diversity (π). Dashed horizontal line indicates the overall average of 0.005. C) Tajima’s D. Dashed horizontal line indicates the overall average of −0.852. D) Linkage disequilibrium Zn scores. Dashed horizontal line indicates the overall average of 0.040.

## Discussion

This is, to our knowledge, the densest genetic linkage map ever created for a conifer species and possible for any tree species. We successfully used this genetic map to anchor 1.7% of the 20 Gbp *P. abies* genome, corresponding to 2.8% of the v1.0 genome assembly (Nystedt et al. 2013), to 12 LGs, constituting the haploid chromosome number (Sax and Sax 1933). The *P. abies* genome has a very large proportion of gene-poor heterochromatin, so while the fraction of the genome that we successfully anchored to the assembly is relatively small, those anchored scaffolds cover 24% of all gene-containing assembly scaffolds and 25% of all partially validated gene models from Nystedt et al. (2013).

The individual LGs from the three component maps (36 LGs from three independent maps) consisted of 648-1,967 probe-markers and 305-1,185 probe-marker bins and, as such, it was not feasible to analyze the maps using an exhaustive ordering algorithm (Mollinari et al. 2009). Instead, we used RECORD (Van Os et al. 2005) with 16 times counting, parallelized over 16 cores and with reordering of markers within 10 marker windows, for each LG to determine the most likely marker order. An heuristic approach, such as RECORD, will undoubtedly introduce some errors in marker ordering (Mollinari et al. 2009), but analyses from simulated data suggested that the average distance between estimated and true marker position is small (< 5 markers) for data sets of similar size to ours (Schiffthaler et al. 2017). However, reliable marker ordering requires robust data and the more genotyping errors and missing data that are present, the harder it will be to determine the true order. This in turn will impact the final size of the map, where both errors in marker order and genotyping results in inflation in the size of the map (Cartwright et al. 2007).

By collecting our 2,000 megagametophytes from what we initially thought were five different ramets of Z4006, we accidentally sampled material from at least three unrelated families. This error stemmed from a mix-up of genotypes due to wrong assignment of ramet ID to the different ramets in the seed orchard. Unfortunately, we were not able to assess which megagametophytes were collected from the different putative ramets since the seed bags were pooled prior to DNA extraction and the sampling errors were not detected until after all sequencing was completed. We used a PCA and hierarchical clustering approach to assign samples into three independent clusters, representing three putative maternal families. We also used PCAs of the putative individual families to verify that these clusters were consistent with offspring derived from a single mother tree (Supplementary, Figure S3). However, we nevertheless cannot completely rule out that a small fraction of samples have been incorrectly assigned to the three families and this would lead to inflated map sizes by introducing an excess of recombination events. Another potential confounding issue is tissue contamination. *P. abies* megagametophytes are very small and are surrounded by a diploid seed coat that needs to be removed prior to DNA extraction. If traces of the diploid seed coat remain in the material used for DNA extractions, the haploid samples will be contaminated with diploid material. To identify and eliminate this possibility, we called sequence variants using a diploid model and any heterozygous SNP calls were subsequently treated as missing data. Samples with a high proportion of heterozygous (>10 %) or missing calls (>20%) were excluded from further analyses to reduce the possibilities of genotyping error due to tissue contamination influencing downstream analyses. We estimated map lengths from 100 rounds of subsampling of 100 random probe-marker bins per component LG and used this to demonstrate that individual maps showed size inflations of 0.15-0.31 cM per probe-marker bin. This inflation is on the same order as the map resolutions for the different clusters and, therefore, indicated an average of ∼1 genotyping error per probe-marker bin or 11-17 genotyping errors per sample.

Both sample‐ and tissue contaminations can influence the accuracy of the genetic map, both with regards to marker order and map size. The smaller family sizes resulting from dividing our original 2,000 samples into three independent families yielded lower resolution of the three component maps. Fortuitously enough, however, this also enabled us to incorporate more markers into the consensus map since different markers were segregating in the different mother trees from which the three families were derived. Furthermore, it also allowed us to evaluate marker ordering across three independently derived maps. Although our consensus map was 70-90% (60-120% for the individual component maps) larger than previously estimated *Picea* maps (3,556 cM vs. 1,889-2,083 cM), it also contained 2-31 times more markers than earlier maps (Pavy et al. 2012; Lind et al. 2014; Pavy et al. 2017). When comparing marker order between our three independent component maps (Cluster 1-3), we found overall high correlations of marker order (0.94-0.99, Supplementary, Figure S8), which is similar to what has previously been observed between estimated and true positions in maps derived from simulated data without genotyping errors but with 20% missing data (Mollinari et al 2009; Schiffthaler et al. 2017). Also, earlier *Picea* maps were all based on diploid F_1_ crosses with even the densest composite map containing only 2,300-2,800 markers per framework map (Table 1 - Pavy et al. 2017), compared to our haploid component maps that contained between 3,924 and 11,479 probe-marker bins each (Table 2).

The comparisons between our haploid consensus map and earlier maps in *Picea* showed an overall high correlation of marker order, which is in line with previous studies suggesting highly conserved synteny within *Picea* and in conifers in general (de Miguel et al. 2015; Pavy et al. 2017). LG I from our haploid consensus map and LG 7 from Lind et al. (2014) showed an inverted order for approximately half of the markers compared (Figure 4). Whether this inversion is due to ordering errors in one of the maps or represents true biological differences between the parents used for the respective maps is, however, not currently known and further investigations are needed to resolve this issue.

A small percentage of the marker comparisons in both the intra‐ and inter-specific maps did not co-align to homologous LGs. Some of these errors likely arose form the repetitive nature of the *P. abies* genome (and conifer genomes in general), where regions with high sequence similarity can often be found interspersed throughout the genome. If the true homologous region between different maps is missing or has been collapsed in the genome assembly due to high sequence similarity, pairwise sequence comparisons may end up assigning homology to regions that are located on different chromosomes. However, it might also be hat these errors represent scaffold assembly errors for scaffolds containing only a single probe-marker or where one region of the scaffold is not captured by the probes, therefore negating evaluation. Approximately 72% of all non-homologues LG comparisons between *P. abies* and *P. glauca* were from multi-markers scaffolds (of which 45% were from probe-markers on split scaffolds in the consensus map (turquoise points in Figure 5). The remaining 28% were comparisons with scaffolds that were only represented by a single probe in the consensus map.

Four percent of the scaffolds containing multiple makers showed a pattern where different markers mapped to different regions, either within or between LGs in the consensus map. This indicates possible errors in scaffolding during the assembly of the v1.0 *P. abies* genome (Nystedt et al. 2013). If this estimate represents the overall picture for the entire assembly, as many as 400,000 of the ∼10 million total scaffolds, and 2,400 of the ∼60,000 gene-containing scaffolds, may suffer from assembly errors. Most worryingly, 2% of the multi-marker scaffolds (100/4,859) contained splits that occurred within a single gene model. It is likely that many of these problematic scaffolds stem from incorrect scaffolding of exons from paralogous genes with a high sequence similarity. Since the *P. abies* genome contains a high proportion of repetitive content, that also includes a large number of pseudo-genes, this is perhaps not surprising. Additional work is needed to disentangle these issues and to resolve any assembly errors. False scaffold joins in a genome assembly are not a unique feature for *P. abies*, rather it appears to be a frequent problem in the assembly process. For instance, dense genetic maps in both *Eucalyptus* and *Crassostrea* have identified and resolved false scaffold joins, thereby improving the genome assemblies in these species (Bartholomé et al. 2015; Hedgecock et al. 2015). Our goal for the *P. abies* genetic map was not only to identify incorrect scaffolding decisions in the v1.0 genome assembly, but to also help improve future iterations of the genome. Long-read sequencing technologies (e.g. Pacific Bioscience or Oxford Nanopore) could be used to resolve these problematic scaffolds and help disentangle the reasons for their ambiguous localization in the genetic map. A future reference genome for *P. a*bies, based on long read technologies will also be able to utilize this genetic map in a much more efficient way since the resulting assembled scaffolds will be substantially longer and would hence enable anchoring a greater fraction of the genome to LGs, ultimately to the point that chromosome-scale assemblies may be achieved.

Our population genetic analyses based on the scaffolds anchored to the consensus map demonstrates the utility of having a dense, accurate genetic map and suggest that the map will facilitate further analyses of genome-wide patterns of variation and selection in *P. abies* in addition to facilitating comparative analyses among spruce species. Assigning even a small fraction of the genome to LGs enabled us to analyze patterns of genetic diversity in approximately a quarter of all predicted genes. This allowed for analyses of broad-scale patterns of variation across the genome and, as the genome assembly is further improved and an even greater proportion of the assembly if physically anchored to the genetic map, will allow for even more fine-scaled analyses of how different evolutionary forces have interacted in shaping patterns of genetic diversity across the *P. abies* genome.

## Acknowledgements

This study was supported by Knut and Alice Wallenberg’s foundation through funding to the Norway spruce genome project. AV was partially supported by a grant from the Stiftelsen Gunnar och Birgitta Nordins fond through the Kungl. Skogs‐ och Lantbruksakademien (KSLA). NRS was supported by the Trees and Crops for the Future (TC4F) project. All computations were performed on resources provided by SciLifeLab and SNIC at the Uppsala Multidisciplinary Center for Advanced Computational Science (UPPMAX) under project b2010042.

## Author contribution

PKI and MRGG conceived the study. AV collected cones and extracted DNA. CB, AV, DS and JB set up bioinformatics pipeline for analyzing sequence capture data. AV and CB performed PCA and identified samples belonging to the three clusters. CB, DS and BS created the genetic maps. CB and PKI performed intra‐ and interspecific map comparisons. CB, XW and PKI performed population genetic analysis. CB performed all remaining analyses and wrote the first draft of the manuscript. NRS contributed to manuscript writing and development of the map construction approach. All authors commented on the manuscript at various stages during the writing.

## Data availability

BatchMap input files for the three clusters, component maps and consensus map files are available from zenodo.org at https://doi.org/10.5281/zenodo.1209841. All scripts needed to recreate the analyses described in the paper are publically available at https://github.com/parkingvarsson/HaploidSpruceMap. Raw sequence data for all samples included in this study are available though the European Nucleotide Archive under accession number PRJEB25757.

